# The Hidden Cost of Fusion: Intrinsic Asymmetry in Vesicle Fusion Restricts Synthetic Cell Growth

**DOI:** 10.1101/2025.09.22.677743

**Authors:** Rafael B. Lira, Rafaela M. Cavalcanti, Karin A. Riske, Rumiana Dimova

## Abstract

Membrane fusion is essential for signaling, cargo delivery, and synthetic cell growth, yet its mechanical consequences remain poorly defined. How fusion-driven membrane growth can be sustained without compromising compartment stability remains an unresolved challenge. Here, we established a minimal reconstituted system where content-loaded small liposomes fuse with single cell-sized giant unilamellar vesicles (GUVs), combining micropipette delivery, electrodeformation, and live imaging. Fusion outcomes were quantified through lipid and content mixing assays, GUV electrodeformation to track area and tension, and phase contrast imaging to monitor leakage. GUVs incorporated lipids and cargo from hundreds of thousands of vesicles at unprecedented efficiency rates, enabling substantial growth. However, accumulation of leaflet asymmetries induced curvature and tension, driving budding, rupture and leakage. Hemifusion amplified these destabilizing effects. Lipid number asymmetries emerge as a dominant mechanical cost of fusion, highlighting how cells may regulate these processes and guiding the design of therapeutic delivery systems and synthetic cells capable of robust and stable growth.

**Significance Statement:** Membrane fusion enables essential biological processes from secretion to cell growth. Using a synthetic system, we show that rapid fusion of small vesicles with model cell-like compartments drives dramatic growth but also introduces leaflet asymmetries that build curvature and tension, limit expansion and compromise membrane stability. By uncovering lipid number asymmetry as a key mechanical cost of fusion, our study explains why cells tightly regulate fusion events and provides principles for designing delivery vehicles and synthetic cells capable of robust, sustainable growth.

## INTRODUCTION

In the course of the cell cycle, living cells must duplicate their components and double their size to prepare for division. A critical aspect of this process is membrane growth, which requires the incorporation of new membrane building blocks (i.e. lipids). In bacteria, lipids are inserted directly into the cytoplasmic membrane(1, 2), while in eukaryotic cells, lipid synthesis occurs primarily at the endoplasmic reticulum (ER) and is followed by redistribution through vesicular trafficking(3, 4), or via secretory vesicles fusion(5). With the aim to reconstitute the cell cycle in vitro, researchers have sought to mimic membrane growth of synthetic lipid compartments using various strategies(6).

In synthetic cell-like compartments, surface area expansion can be accomplished by incorporation of lipids or other amphiphiles through three main strategies: (i) insertion of molecules or micelle supplied in the external medium, (ii) direct local *de novo* synthesis of lipids, or (iii) fusion of vesicles. In the first route, amphiphilic molecules such as detergents, fatty acids or lysolipids, are externally supplied, often in the form of micelles, driving measurable membrane growth(7). However, this is often at the cost of membrane integrity due to their solubilizing effects. Furthermore, this strategy has mostly been demonstrated using population of small vesicles and lipid aggregates that elongate into tubules under external shear(8), limiting its relevance for cell-sized vesicles(9). While spontaneous incorporation of fatty acid might have played a role in early protocells(10), modern cellular membranes are composed of more stable phospholipids(11, 12).

The second route, direct lipid synthesis, more closely reflects the situation in living cells. However, the reconstitution of the required enzymatic machinery (including enzymes, co-factors, and precursors) is technically demanding and prone to variability. Encapsulation of these components often leads to large vesicle-to-vesicle variations(13, 14), and even in successful cases, synthesized lipids result only in minimal membrane growth(15, 16). Furthermore, lipid synthesis is slow, requiring timescales ranging from many minutes(15) to hours(17).

In eukaryotes, vesicular intracellular trafficking through tightly regulated cycles of membrane fission and fusion enables continuous exchange of lipids from their production site to other intracellular organelles(18, 19). Fusion of vesicles contribute to the dynamic regulation of membrane area(20) and surface tension(21) while facilitating mass transfer of lipid and aqueous content between organelles(22). Inspired by this biological framework, membrane fusion has been explored as a strategy to promote growth in synthetic cell systems. However, achieving robust and controllable growth of membrane-bound compartments via vesicle fusion (particularly with cell-sized vesicles) remains a significant challenge.

Few studies have demonstrated growth via fusion in synthetic systems, and even those have shown only modest membrane area increase, whether using lipid-(23) or polymer-based (24) compartments. These increases are very far from the minimal requirement for functional growth, such as area doubling (here, we exclude studies involving fusion between two preformed cell-sized vesicles(25, 26), which do not recapitulate the asymmetry and directionality required for synthetic cell growth). Furthermore, it remains unclear whether repeated fusion events trigger potential structural alterations of the target compartment membrane, an essential prerequisite for developing stable, sustainable and functional synthetic cells. While there is evidence that membranes can become leaky under certain fusogenic conditions(27–31), no current system has demonstrated substantial, non-destructive membrane growth through fusion.

To develop lipid-based vesicular compartments capable of controlled growth (and ultimately division), fusion strategies must meet several criteria: they must be highly efficient, fast, and gentle enough to preserve the integrity and functionality of the compartmental membrane. Fusion of membranes made of cell-compatible building blocks not only advances the design of synthetic cells, but also gives valuable insights into the working principles of membrane fusion and area regulation in cells.

Cationic large unilamellar vesicles (LUVs) can fuse very efficiently with both living cells(32) and cell-sized phospholipid giant unilamellar vesicles (GUVs) (33). The fusion process can be finely-tuned by adjusting biophysical parameters such as membrane charge density(33) and fluidity(34). Unlike protein-mediated fusion, which relies on complex biochemical reconstitution of proteoliposomes(35), fusion with charged LUVs involves far simpler preparation and is significantly more fusogenic (Lira and Witkowska, in preparation), making it an attractive strategy for fusion-mediated growth of synthetic membrane compartments.

In this work, we fuse cationic LUVs with GUVs as a route to achieve unprecedent growth of cell-like synthetic cell compartments. Although membrane fusion has been widely studied, only few studies have demonstrated considerable membrane area increase, and even fewer have quantified it. To quantify fusion efficiency, membrane area growth, and any resulting morphological or structural changes, we combined single-vesicle spectroscopy, micromanipulation and real-time microscopy. The use of fluorescence resonance energy transfer (FRET) and content mixing assays helped us assess fusion efficiency of LUVs locally injected in the vicinity of GUVs. We measured the resulting GUV area increase and investigated how fusion of a large number of inherently asymmetric LUVs affects GUV morphology and mechanics, with particular attention to the resulting GUV transmembrane lipid asymmetry. Specifically, we examined how vesicle size, fusion frequency, and lipid leaflet imbalances influence the capacity of GUVs to accommodate new membrane material. Our findings underscore the critical role of lipid number asymmetries between bilayer leaflets in shaping the growth, stability, and mechanical response of synthetic membrane compartments. These insights offer a step toward designing vesicle systems capable of sustained growth and functional behavior, and may illuminate fundamental aspects of membrane dynamics relevant to both synthetic biology and cellular membrane homeostasis.

## RESULTS

To drive GUV growth via fusion, we prepared cationic LUVs from mixtures of the positively charged lipid dioleoyl trimethylammonium propane (DOTAP), the cone-shaped lipid dioleoyl phosphoethanolamine (DOPE) and a small fraction of the FRET acceptor dye lissamine-rhodamine-labelled PE lipid (DPPE-Rh) at 1:1:0.1 molar ratio. DOTAP alone is sufficient to mediate fusion of cationic liposomes with GUVs, whereas the addition of DOPE further enhances fusogenicity(36).

The LUVs were incubated with neutral GUVs composed of pure palmitoyl-oleoyl phosphatidylcholine (POPC), or with negatively charged GUVs composed of mixtures of POPC and palmitoyl-oleoyl phosphatidylglycerol (POPG) at molar ratios of 5:5 unless stated otherwise. All GUV compositions included trace amounts of the FRET donor DPPE-NBD [1,2-dipalmitoyl-sn-glycero-3-phosphoethanolamine-N-(7-nitro-2-1,3-benzoxadiazol-4-yl) (ammonium)] in the membrane.

### Quantification of membrane fusion efficiency

Here, we employed three independent methods to quantify the increase in GUV membrane area resulting from the fusion of LUVs: (i) a lipid-mixing FRET assay, (ii) a content-mixing assay based on encapsulated dye, and (iii) direct quantification of area increase using GUV electrodeformation. Notably, the latter two approaches are applied here for the first time to quantitatively evaluate fusion at the single-vesicle level. By integrating quantitative imaging with micromanipulation, we were able to assess not only the number of LUVs fused per GUV, and, thus, the amount of material delivered to the GUV membrane and lumen, but also the resulting changes in GUV composition, leaflet asymmetry, membrane integrity, and area – all in a single-vesicle, self-contained experiment.

In method (i), fusogenic cationic LUVs containing the FRET acceptor probe DPPE-Rh fused with GUVs containing the FRET donor molecule DPPE-NBD. The FRET efficiency (*E*_*FRET*_) is measured directly on the GUV membrane and defined as *E*_*FRET*_ = *I*_*Rh*_⁄(*I*_*NBD*_ + *I*_*Rh*_), where *I*_*Rh*_ is the fluorescence intensity in the FRET channel (excitation of NBD and emission of Rh due to FRET) and *I*_*NBD*_ is the fluorescence intensity of the donor probe (direct excitation and emission of NBD). Fusion efficiency can be assessed using confocal fluorescence microscopy based on FRET between membrane-incorporated donor (DPPE-NBD) and acceptor (DPPE-Rh) dyes. We monitored FRET changes via ratiometric imaging, which offers higher temporal resolution than lifetime-based measurements(33, 34, 36, 37), enabling us to resolve fast fusion kinetics. Figure S1 shows how we measure membrane intensity. In the absence of fusion, GUVs lacking the FRET acceptor exhibit *E*_*FRET*_ = 0. To avoid confounding effects from reduced *E*_*FRET*_ caused by LUV docking, measurements were restricted to regions free of docked vesicles. Fusion ideally results in the transfer of the acceptor (DPPE-Rh) to the GUV membrane, yielding *E*_*FRET*_ > 0. Thus, the extent of FRET, and hence the relative donor-acceptor ratio in the GUV, can serve as a quantitative proxy for the number of LUVs that have fused with a given GUV. To calibrate the relationship between FRET efficiency and lipid mixing, we constructed a FRET curve using GUVs containing different molar fractions of DPPE-Rh (0–5 mol%) while maintaining a constant amount of DPPE-NBD (0.5 mol%) (Figure S2A,B). This calibration emulates the progressive incorporation of acceptor lipid during fusion and enables us to estimate the fraction of lipids (*X*_*fuso*_) transferred from LUVs to GUVs during fusion.

For method (ii), we assessed content mixing using the water-soluble dye Atto, which was encapsulated inside LUVs. To quantify dye transfer into GUVs upon fusion, we first constructed a fluorescence calibration curve of free Atto at known concentrations in solution (Figure S2C). Content mixing serves as an independent fusion readout and helps distinguish full fusion (where both lipid and aqueous contents are exchanged) from partial intermediates such as docking or hemifusion (which can produce FRET signal). An important advantage of these single-vesicle fluorescence microscopy methods over bulk fluorimetry is their ability to simultaneously reveal morphological changes, such as area expansion, membrane fluctuations, and budding, which also serve as sensitive indicators of fusion extent and outcome.

We performed end-point measurements of lipid and content mixing on individual GUVs after 10-15 minutes of incubation with increasing concentrations of LUVs, a duration previously shown to be sufficient for fusion based on isothermal titration calorimetry(38), LUV-GUV mixing in microfluidic traps(33) and local LUV microinjection (see below). Representative confocal images (Figure 1A) show that both lipid and content mixing in negatively charged GUVs progressively increase with LUV concentration. Interestingly, the GUV morphology also undergoes noticeable changes depending on the LUV concentration. The fusion experiments were performed in mildly hypotonic solutions to ensure that changes in vesicle area/volume are solely a result of fusion (i.e. growth).

**Figure 1.**
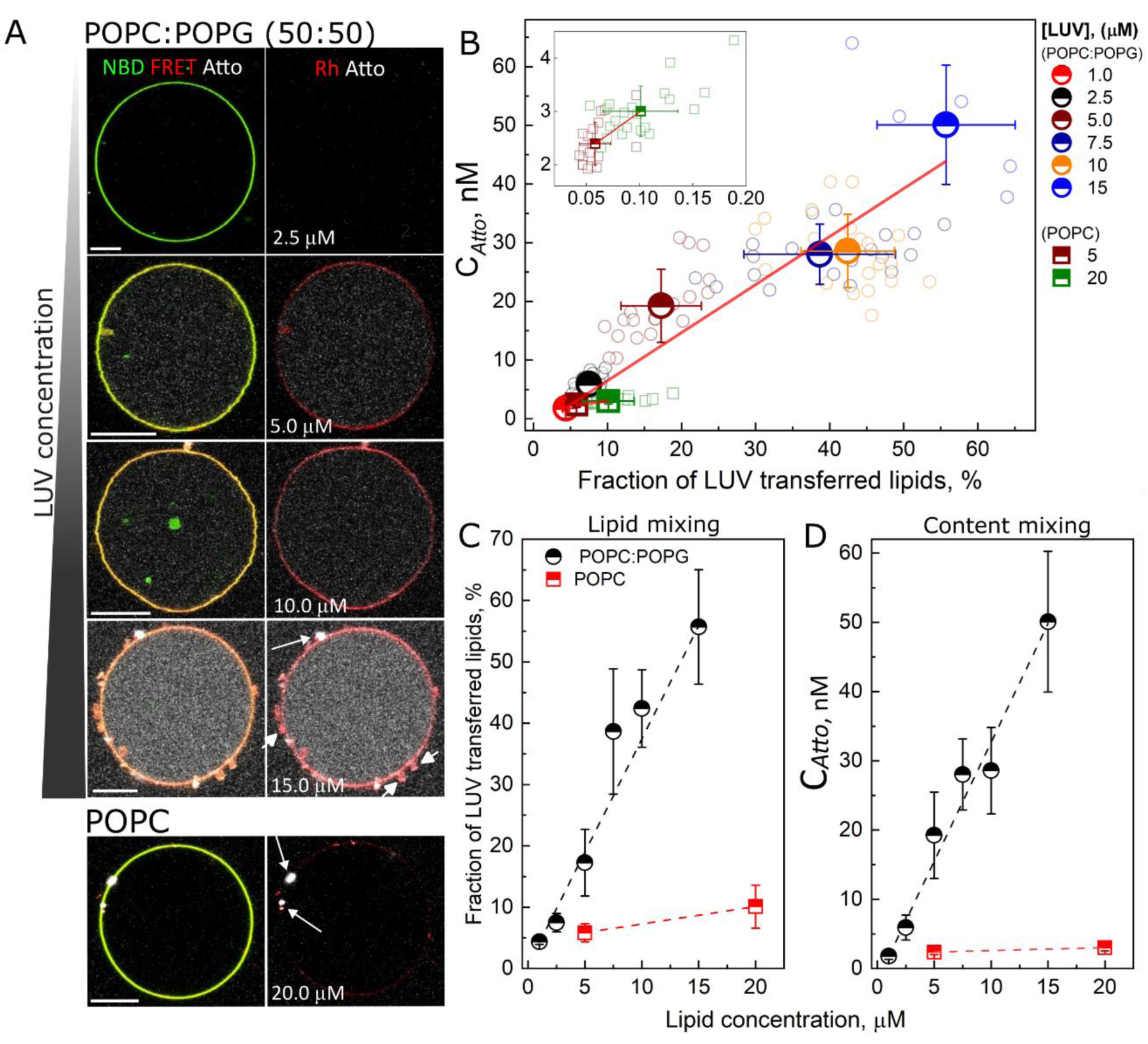
Quantification of fusion efficiency from bulk incubation of GUVs with fusogenic LUV solutions via simultaneous detection of lipid and content mixing. (A) Snapshots of negatively-charged POPC:POPG GUVs (1:1, molar ratio) incubated with increasing concentrations of LUVs. A neutral POPC GUV incubated with LUVs is also shown at the bottom. Left column: overlay of FRET donor channel (NBD, green), acceptor channel (FRET, red) and encapsulated content marker (Atto emission, white). Right column: overlay of DPPE-Rh and Atto signals (both initially located in the fusogenic LUVs) upon direct excitation, highlighting LUV docking as bright white puncta on the GUV surface (arrows). Atto was encapsulated in the LUVs at a concentration of 20 μM. The very faint signal in the Rh channel (red) of the POPC vesicle is due to low fusion efficiency. (B) Atto concentration in the GUV lumen (indicative of content mixing) as a function of the fraction of GUV lipids originating from LUV fusion (*X*_*fuso*_) for both POPC and POPC:POPG GUVs. Individual data points (open symbols), means (half-filled symbols), and standard deviations are shown. Inset: magnified view of the data for POPC GUVs. (C, D) fraction of lipids in the GUVs transferred from the LUVs and Atto concentration (*C*_*Atto*_) as functions of LUV concentration (in terms of lipid concentration). Means and standard deviation from the measurements shown in B. Linear fits are shown for each dataset. Scale bars: 10 μm.

At low LUV concentrations, both FRET and Atto signals indicate successful fusion events (lipid and content mixing), and the GUVs remain spherical (Figure 1A, 5 μM lipids). At intermediate concentrations, fusion becomes more pronounced: both lipid and content mixing signals increase, the GUVs are no longer spherical but exhibit visible membrane fluctuations, indicative of excess area (Figure 1A, 10 μM lipids). At high LUV concentrations, GUVs return to a more spherical appearance, but now contain numerous small membrane buds (arrows in Figure 1A, (Figure 1A, 15 μM lipids) and no longer show membrane fluctuations, suggesting that excess area has been compartmentalized into protrusions. Fusion reactions at higher LUV concentrations were not pursued because they often caused GUV rupture, characterized by the formation of large pores, vesicle bursting, and transformation into tubular structures(36). As discussed further below, this behavior suggests that increased spontaneous curvature effects lead to a reduction in both the number and size of GUVs as the LUV concentration rises.

Figure S3 presents measurements of Atto intensity inside the GUV lumen as a function of *E*_*FRET*_, showing that in negatively charged GUVs, lipid and content mixing occur in a coupled manner. Using the calibration data (Figure S2) and the raw data in Figure S3, Figure 1B displays the calculated Atto concentration in the GUV interior (*C*_*Atto*_) as a function of *X*_*fuso*_, the estimated fraction of GUV lipids that originate from LUVs via fusion (see Methods for derivation). Under the tested conditions, fusion of 15 μM LUVs (lipid concentration) results in the delivery of up to 50 nM Atto (initially encapsulated in the LUVs at 20 μM concentration), with more than 50 mol% of the GUV membrane lipids originating from the LUVs. The data confirm that LUVs fuse extensively with GUVs, significantly increasing their membrane area and altering their lipid composition. Figures 1C and 1D show that both *X*_*fuso*_ and *C*_*Atto*_ increase linearly with LUV concentration across a large number of GUVs.

We also examined whether the LUVs fuse as efficiently with electrically neutral GUVs. Incubation of LUVs with neutral POPC GUVs resulted in detectable fusion as indicated by an increase in *E*_*FRET*_ (reflecting lipid mixing) with raising LUV concentrations (see also Figure S3). However, for comparable LUV concentrations, lipid mixing was much lower in neutral than in negative GUVs (Figure 1B,C). More importantly, changes in *E*_*FRET*_ were only modestly accompanied by content mixing signal, even at higher LUV concentrations, suggesting that fusion with neutral GUVs predominantly results in docking and hemifusion – a finding consistent with previous lipid quenching experiments(33). At high LUV concentrations, we still detect a small amount of Atto in the GUV lumen, indicating that some content mixing does occur.

### Real-time fusion

The bulk experiments described above allowed us to study LUV fusion with GUVs under controlled and increasing LUV concentrations. While robust, this approach yields only end-point measurements and lacks temporal resolution and morphological context. To capture the kinetics and morphological changes of the GUV membrane during fusion, we performed real-time experiments by microinjecting LUVs near individual GUVs and imaging their response dynamically.

The LUVs were microinjected approximately 20–50 µm from the GUV using a micropipette. Upon exiting the injection pipette, LUVs are rapidly diluted in the surrounding medium. Thus, we used a high LUV concentration (420 μM lipid concentration) to load the micropipettes. Imaging was performed using multicolor confocal microscopy combined with phase-contrast to simultaneously visualize lipid mixing, content mixing, and morphological changes. GUVs were prepared in sucrose and dispersed in glucose, rendering them optically distinct (appearing darker and with a bright halo) under phase contrast microscopy due to differences in the refractive index. Loss of optical contrast served as an indicator of increased membrane permeability and solute exchange, often associated with pore formation. Figure 2A shows a configuration of the experiment and the Supporting Information (SI) and Figure S1 detail the quantification method.

**Figure 2.**
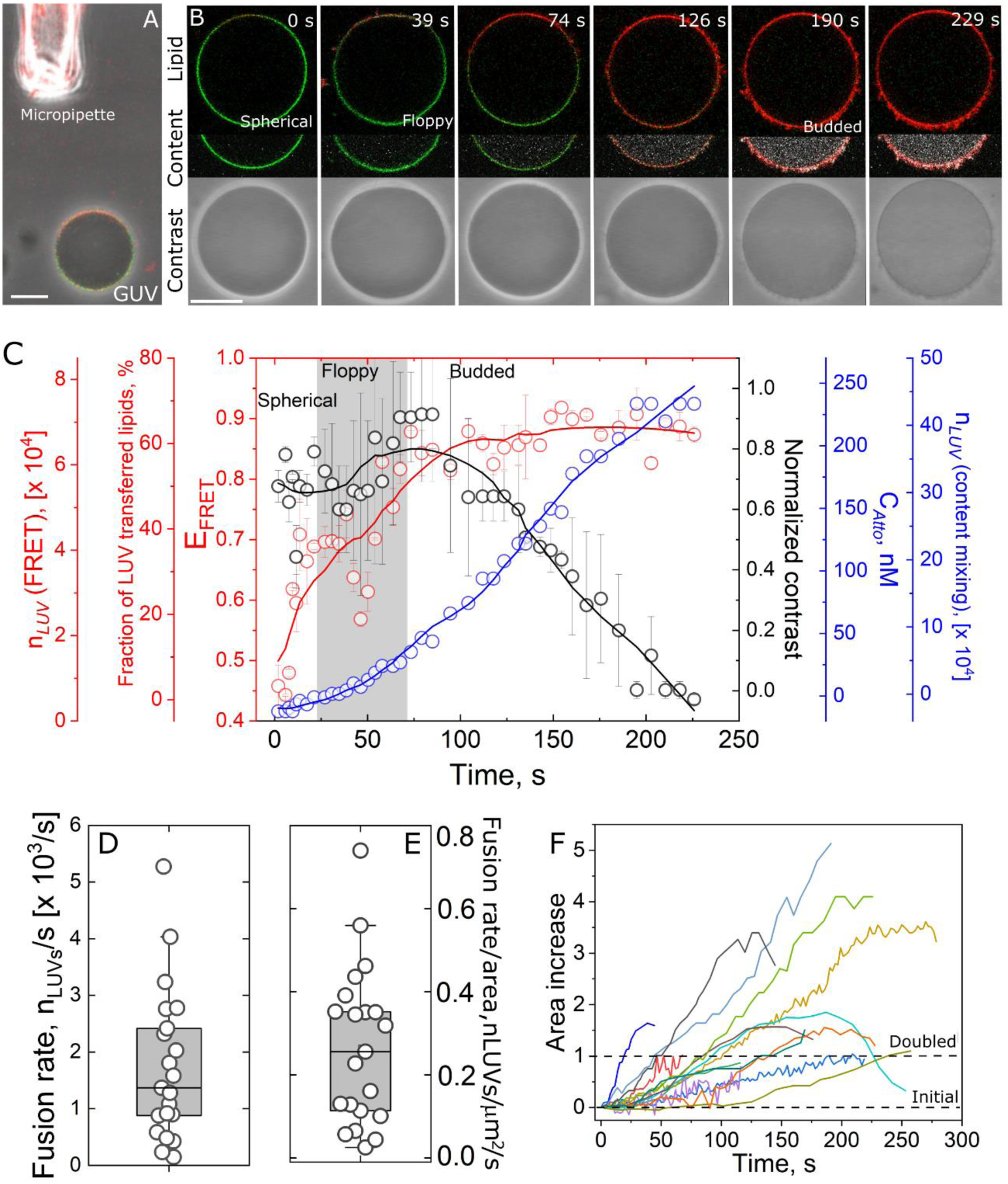
Real-time monitoring of LUV-GUV fusion via micropipette injection. (A) Confocal/phase-contrast overlay of a GUV during local injection of LUVs (420 μM lipid concentration) from a micropipette. Fusion begins immediately on the membrane side facing the pipette, indicated by red fluorescence. Scale bar: 10 µm. (B) Time-lapse sequence of a GUV during fusion. Top row: lipid mixing visualized by overlaying direct emission from the donor NBD (green, in the GUV) and the acceptor Rh (red, in the LUVs); middle row: content mixing (including Atto signal, grayscale); bottom row: phase-contrast imaging. Scale bar: 20 µm. (C) Quantitative analysis of *E*_*FRET*_ (lipid mixing), fraction of membrane lipids originating from LUVs, *C*_*Atto*_ (lumenal dye concentration), *n*_*LUV*_ (number of fused LUVs), and normalized optical contrast for the GUV shown in B. Errors are from two independent measurements on the same GUV. Circles show measured values; lines are a guide to the eye. Representative GUV morphologies characteristic for the regions indicated in the graph are shown in panel B. (D, E) Box plots of calculated fusion rates: absolute (D) and normalized by GUV area (E). Data points are measurements on individual GUVs. (F) Estimated fractional increase in GUV membrane area, Δ*A*/*A*_0_, based on content mixing. The dashed lines represent the initial GUV area before fusion and its estimated doubling. The GUVs (POPC:POPG, 5:5 molar ratio and labelled with 0.5 mol% NBD) were prepared in 200 mM sucrose and dispersed in 185 mM glucose.

Immediately upon injection, *E*_*FRET*_ increases locally at the GUV side facing the micropipette (Figure 2A,B). This signal later spreads due to lateral lipid diffusion and further fusion events across other parts of the GUV. Concurrently, Atto fluorescence in the lumen and GUV morphology change progressively, capturing the entire fusion trajectory (Figure 2B and Movie S1). The initially spherical GUV gains membrane area and volume, resulting in pronounced shape fluctuations (e.g., floppy membranes at 39 s). Continued fusion triggers budding transitions, which store excess membrane area (e.g., at 126 s). Eventually, the GUV becomes permeable, as indicated by progressive loss of phase contrast (>190 s), consistent with submicron pore formation. Notably, the onset of contrast loss occurs stochastically but reproducibly across different GUVs. Pore formation leads to delayed leakage of Atto (compared to sucrose/glucose), due to its lower diffusion rate and continued supply (see Figure 4).

Quantitative analysis in Figure 2C shows the broad spectrum of information that can be obtained from the experiment. The values of *E*_*FRET*_ rise to saturation levels. Note that the measurements were taken from the membrane region facing the pipette, leading to precise but likely overestimated global values in the beginning of the experiment. Furthermore, rather than complete lipid replacement and saturation of fusion, the saturation of *E*_*FRET*_ reflects measurement limits introduced by the definition of this quantity (for *I*_*Rh*_ ≫ *I*_*NBD*_, *E*_*FRET*_ approaches 1, regardless of the absolute amount of lipid mixing). Using the calibration curve in Figure S1, we estimated that 60,000 LUVs fuse with a single GUV during the experiment. This corresponds to a substantial change in local membrane composition: by the end of the measurement, approximately 60% of the GUV membrane lipids originate from the LUVs.

A complementary estimate can be derived from the content-mixing signal. Because Atto diffuses rapidly in the GUV lumen (diffusion coefficient ∼400 μm²/s(39)), the resulting fluorescence signal is spatially homogeneous and provides a reliable bulk measure of lumenal content transfer. When LUVs encapsulating 20 μM Atto fuse with GUVs, we measure a final Atto concentration in the GUV lumen of approximately 250 nM. This corresponds to ∼500,000 LUVs fusing with a single GUV, yielding a high fusion efficiency (*X*_*fuso*_ or around 75%). To the best of our knowledge, this represents the highest level of fusion efficiency reported for any membrane fusion system, although quantitative reports of fused LUVs (*n*_*LUV*_) are rare in the literature(40).

As can be expected, the number of fusion events (*n*_*LUV*_) scales with GUV size – larger GUVs accommodate more fusing LUVs (Figure S4 A). Interestingly, we observed that at later stages of the fusion process, GUVs become permeable. In the example shown in Figure 2A-C, contrast loss begins around 130 seconds and continues until the enhanced optical contrast conferred by the sugar asymmetry is lost. This indicates that the membrane can support a substantial number of fusion events while remaining intact, enabling the fusion of tens of thousands of LUVs (corresponding to approximately a doubling of membrane area) before its integrity is compromised. Beyond a certain threshold, however, membrane pores form and leakage is sustained for at least 1– 2 minutes (once the contrast is lost, we do not have a readout for leakage). While the onset and kinetics of these transitions vary depending on local LUV concentration and GUV size, the sequence of events is qualitatively reproducible for all GUVs.

Because *n*_*LUV*_ is measured dynamically, we can calculate the fusion rate directly. From this point onward, we focus on the content mixing data, as the FRET signal saturates at high *n*_*LUV*_, and does not account for donor dilution caused by membrane area expansion. We determined fusion kinetics using the linear phase of the content mixing signal (Figure S4B). Figure 2D shows that, on average, ∼1,500 LUVs fuse with a GUV per second, with peak rates reaching up to ∼5,000 LUVs per second in some cases. When normalized to the GUV membrane area, this corresponds to a fusion rate of approximately 1 LUV per 3 µm² per second (Figure 2E). These data not only reveal a remarkably high extent of fusion, but also demonstrate that fusion occurs at extraordinary speed. To our knowledge, such rapid and extensive fusion has not been reported in other GUV fusion systems, making this result particularly noteworthy.

Given this high fusion efficiency, we next estimated the resulting increase in GUV membrane area due to LUV fusion (see approach in SI). This calculation allows us to quantify the growth of the GUV compartment via fusion. A fractional area increase of 0 corresponds to no growth, while a value of 1 (100%) reflects a doubling of the membrane area. Figure 2F presents the fractional area increase for 21 individual GUVs. Most GUVs display at least a two-fold increase in area, with some showing substantially greater expansion. It is worth noting that particularly high values may be overestimates, as GUV volume tends to decrease at high *n*_*LUV*_ values (Figure S5), likely due to budding, leakage and increased membrane tension. This could result in an elevated lumenal Atto concentration (*C*_*Atto*_), which affects area estimation. Additionally, GUVs that develop pores experience Atto leakage, leading to an apparent reduction in calculated area gain at later time points.

In summary, the exceptionally high level of LUV fusion not only increases GUV membrane area substantially, but also underscores the potential of this system to support growth and area doubling. The latter represent key prerequisites for constructing synthetic compartments capable of division and self-replication.

### Effective increase in GUV area is only partially available for compartment growth

Membrane fusion inherently increases the surface area of the involved compartments, such as cells, organelles, or GUVs. However, this increase is rarely directly measured or detected, with only a few exceptions. Previous studies have mostly relied on indirect assessment, employing approaches such as dynamic light scattering of small liposomes(38, 41), electron microscopy(27, 42, 43), or detection of modest overall GUV size increases induced by osmotic swelling combined with fusion(23) and low-efficiency DNA-mediated fusion(44). These techniques typically depend on ensemble measurements, which obscure heterogeneities within samples. Additionally, some methods involve sample destruction (e.g., electron microscopy), while others lack sufficient efficiency to generate measurable growth. Our system overcomes these limitations by enabling direct, high-resolution, real-time measurement of membrane area changes at the single-vesicle level.

To determine whether this gained membrane area is functionally available for compartment growth rather than accumulated in defects, we applied a weak alternating current (AC) electric field to mechanically deform the GUVs and pull out the deformations (45, 46). Electrodeformation is a well-established technique to quantify membrane area changes and mechanical properties(46, 47), having been previously used to assess area expansion upon insertion of amphiphilic molecules(7, 48), lipid photooxidation(49) and to quantify GUV excess area(50)

In these experiments, GUVs were slightly inflated using mildly hypotonic conditions to minimize pre-existing excess area, ensuring that observed deformations reflect fusion-mediated area gain. Electrodeformation was performed simultaneously with lipid and content mixing measurements, providing complementary and independent methods to assess are increase.

When GUVs were exposed to increasing bulk concentrations of LUVs, we observed clear and quantifiable deformations up to ∼ 10 µM LUV lipid concentration (Figure S6). At this concentration, on average, 3,000 LUVs fuse to single GUVs, resulting in substantial membrane area gain. At 15 µM lipid concentration, deformation was still observed but began to coexist with membrane budding, limiting the use of electrodeformation to accurately quantify area. At even higher concentrations, the GUVs no longer deformed in response to the AC field. These results suggest that although fusion delivers substantial new membrane material, not all of it remains available as free surface area of the GUV compartment. Instead, a portion is stored in membrane structures such as buds, which impose mechanical tension on the GUV and limit further deformation.

To gain dynamic insight into this transition, we performed real-time imaging of GUVs undergoing LUV fusion under an AC field (Figure 3A). Based on the above results, we hypothesized the following sequence: (i) the GUV is initially spherical and undeformed (i.e. tense by the slight osmotic inflation), (ii) as LUVs fuse, membrane area increases, allowing the GUV to deform under the field, (iii) further fusion drives maximum deformation, and (iv) the GUV undergoes a budding transition that counteracts further deformation.

**Figure 3.**
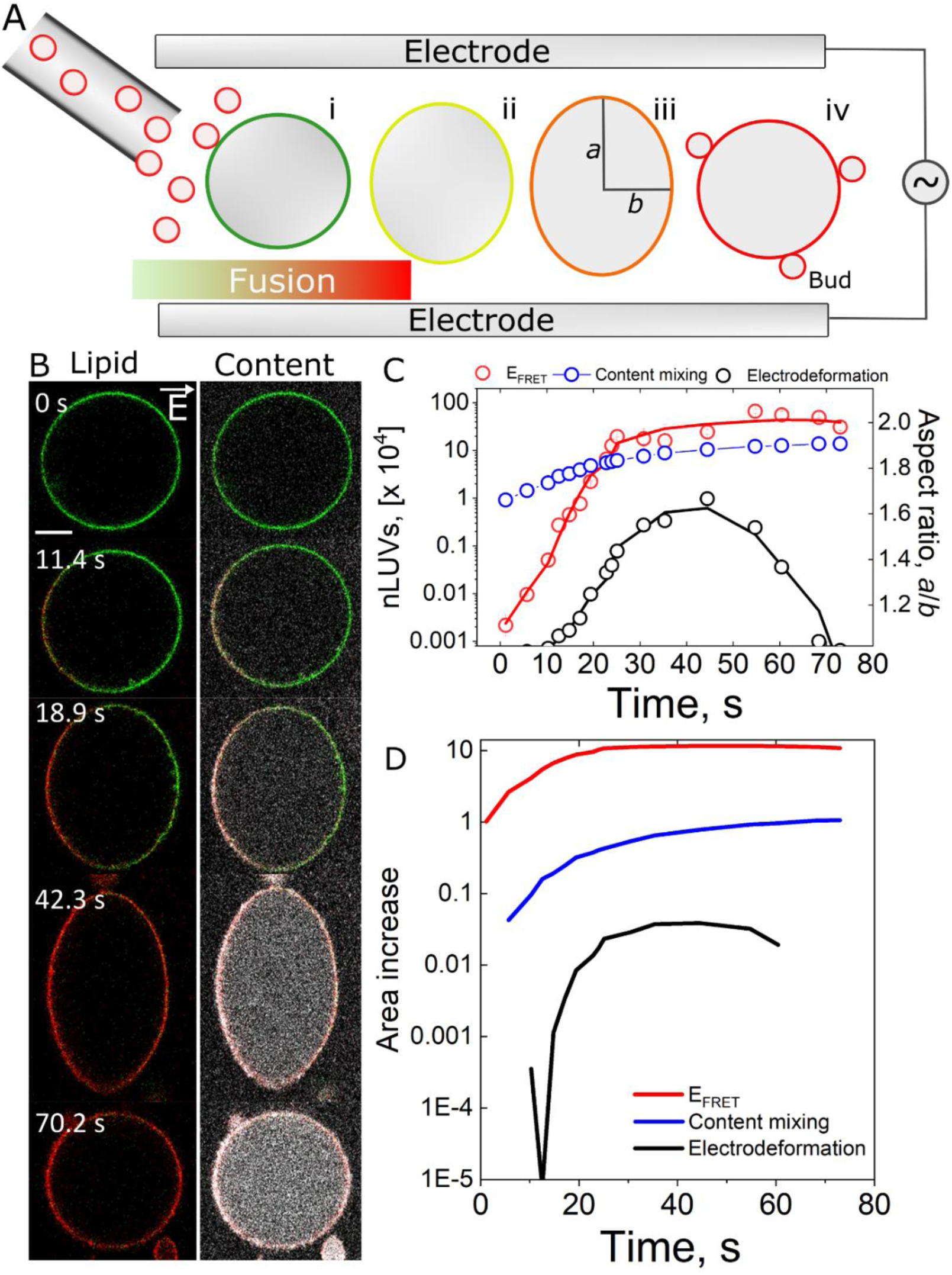
Real-time electrodeformation of GUVs during membrane fusion for assessing area changes. (A) Schematic of the electrodeformation setup. GUVs are placed between two parallel electrodes spaced 500 µm apart, and exposed to an AC field. Fusogenic LUVs (red) are delivered near the GUVs using a micropipette and reach the GUV membrane by diffusion. The electric field is applied continuously throughout the experiment. Fusion is monitored in real time via lipid mixing (FRET) and content mixing (Atto dye); morphological changes such as budding due to increased spontaneous curvature are also shown. (B) Time-lapse confocal microscopy images from a typical experiment. The AC field was set to 10 V_pp_ at 500 Hz. Left column: lipid mixing visualized by overlaying direct emissions from the FRET donor (NBD in the GUV) and acceptor (Rh in the LUVs). Right column: content mixing visualized via Atto dye (grayscale). Scale bar: 10 µm. (C) Quantification of GUV fusion and deformation. Plotted are the GUV aspect ratio (*a*/*b*), the estimated number of fusing LUVs (*n*_*LUV*_) from both lipid (FRET) and content mixing assays, and the effective area increase derived from electrodeformation. (D) Comparison of GUV area increase, *α* = Δ*A*/*A*_0_, estimated independently from FRET, content mixing, and deformation analysis. The GUVs were prepared in 200 mM sucrose containing 1 mM NaCl inside and mildly inflated by dilution in a slightly hypotonic 185 mM glucose solution to ensure no excess area prior to fusion.

Figure 3B shows time-lapse images of a typical experiment that supports this sequence. Initially, the mildly inflated GUV is spherical and exhibits no deformation (at the onset of injection, t = 0 s,). As fusion progresses, evident from increases in FRET and Atto dye transfer, the GUV deforms (t = 11.4–18.9 s) and continues to elongate (t = 42.3 s). Eventually, as more LUVs fuse, the GUV returns back to a spherical shape and forms visible membrane buds. The full process is shown in Movie S2.

Figure 3C displays the evolution of the GUV aspect ratio (*a*/*b*) over time for the vesicle shown in Figure 3B, along with the estimated number of fused LUVs (*n*_*LUV*_) derived from lipid mixing (*E*_*FRET*_), content mixing (*C*_*Atto*_), and electrodeformation (*a*/*b*). The apparent decrease in *n*_*LUV*_ based on electrodeformation at *t* > 40 s does not reflect actual loss of fusion events, but rather marks the transition point at which membrane tension exceeds the deforming force from the electric field. At this point, the values of *n*_*LUV*_ estimated from lipid and content mixing reach ∼10⁵–10⁶, while the estimate from electrodeformation remains significantly lower (∼10³-10^4^).

To further compare these methods, Figure 3D shows the calculated membrane area increase based on each approach (see SI for calculation details). The relative area increase is expressed as *α* = Δ*A*/*A*_0_, where *A*_0_ represents the initial GUV area prior to the onset of fusion; note that *α* = 1 corresponds to doubling the GUV area. The FRET-based estimate of *α* is markedly overestimated, primarily because measurements are taken from the membrane region facing the pipette (where lipid accumulation is most concentrated), leading to artificially inflated values. In contrast, the area increase calculated from content mixing reflects a more accurate and spatially averaged measurement, revealing a doubling of membrane area.

However, the area mechanically available for deformation, as determined from maximum electrodeformation, accounts for only ∼ 4% increase in this particular GUV. This trend is consistent across multiple GUVs (see Figure S7), where the average area gain estimated from content mixing was 2.3 ± 1.1-fold, or 130% (n = 8), while the maximum deformation-derived values for α is 2.2±0.8%, ranged from 1.4 – 4.0%. Obviously, the extent of observable deformation depends on the method employed. Electrodeformation using AC fields operates in a relatively low-tension regime, thus reporting only a small fraction of the total area gained.

Interestingly, the area increase measured by electrodeformation exhibits a slower response than that inferred from content mixing (see Figures 3D and S7). This delay is likely due to the initial accumulation of fused membrane material on the side of the GUV facing the pipette; for this excess area to result in measurable GUV deformation under the applied field, it must first redistribute across the vesicle surface. The degree of delay and deformation also appears to depend on GUV size, as reflected in the variability shown in Figure S7. Once the maximum deformation is reached, much of the additional membrane area becomes stored in highly curved structures, notably membrane buds.

### Fusion creates lipid number asymmetry, induces spontaneous curvature, and builds membrane tension

The observed decline in the apparent area gain as measured by electrodeformation at later stages of fusion reflects the dynamic balance between electrically induced membrane tension and fusion-driven remodeling. When an electric field is applied, the GUV adopts a prolate ellipsoidal shape due to increased membrane tension. This electric-field-induced tension can be estimated from the applied field strength and vesicle morphology, specifically, from the mean curvatures at the poles and equator(51).

As fusion proceeds, the GUV membrane gains area and initially deforms more. However, the incorporation of large numbers of highly curved LUVs introduces increasing asymmetry between the inner and outer leaflets of the GUV. Due to their small size (*R*_*LUV*_ ≈ 60 nm), LUVs are inherently asymmetric, containing more lipids in their outer leaflet. This asymmetry is transferred to the GUV during fusion and accumulates over time, leading to an increase in spontaneous curvature – the membrane’s inherent tendency to curve in the absence of external forces(52). The buildup of spontaneous curvature gives rise to budding and introduces a “spontaneous tension”(52), which resists further deformation and promotes a return to a spherical shape.

This spontaneous tension, given by *σ*_*m*_ = 2*km*^2^ (52), (where *κ* is the bending rigidity and *m* the spontaneous curvature), can eventually exceed the electric-field-induced tension. For the vesicle in Figure 3B, the electric tension at the point of maximal deformation (∼40s) is estimated to be ∼ 0.5 mN/m, setting a lower bound for *σ*_*m*_ at this maximal deformation. The corresponding spontaneous curvature is approximately *m* ≈ 1/22 nm⁻¹. For small spherical buds, the relation *m* = 2/*D*_*bud*_ (52, 53) gives a characteristic bud diameter of approximately 44 nm, which is well below the diffraction limit, consistent with experimental observations. In principle, using a feedback-controlled increase in the electric field to reopen membrane buds could enable real-time tracking of changes in spontaneous curvature. However, such an approach is experimentally demanding, as the applied field would need to be carefully tuned to each vesicle’s size and deformation profile and the inhomogenous surface charge might play a role. As we discuss below, time-resolved curvature changes can still be inferred from bud morphology observed in the absence of the field.

In the field-free case (Figure 2B), budding is driven solely by the fusion-mediated accumulation of membrane asymmetry. At low fusion levels, these effects are minimal, but with increasing number of fusing LUVs, the GUVs transition to a budded state. In fact, GUVs initially exhibiting inward tubes (indicative of negative spontaneous curvature) can reverse asymmetry, whereby the tubes are suppressed and outward buds form with increasing fusion, confirming the role of LUV fusion in generating positive curvature (Figure S8, Movie S4). In some cases, the presence of tubes appears to buffer the membrane from poration and rupture, suggesting a protective role through area storage.

This budding behavior (in the absence of electric field) allows an estimate of *m* from bud diameter measurements. In the early fusion stages, the asymmetry (i.e. spontaneous curvature) is low and the buds are large (*D*_*bud*_ = 2/*m*). With increasing asymmetry, bud size decreases and resolving small buds is challenging due to their dynamics and the limited optical resolution. In rare cases, buds are stable and in focus throughout the fusion process, enabling dynamic measurements. Figure 4A shows such an example, with bud size evolution tracked in Figure 4B,C. Here, spontaneous curvature increases from <1 μm⁻¹ to ∼7 μm⁻¹ i.e. ∼1/(150 nm). This corresponds to a rise in *σ*_*m*_ from near zero to ∼0.08 mN/m, consistently lower than the tension assessed from electrodeformation experiments. Although significant, this value remains well below the lysis tension of fluid membranes, which around ∼1-5 mN/m (54, 55), but as fusion proceeds, can exceed it, consistent with the observed GUV leakage (Figures 4A and S5) and occasional rupture.

**Figure 4.**
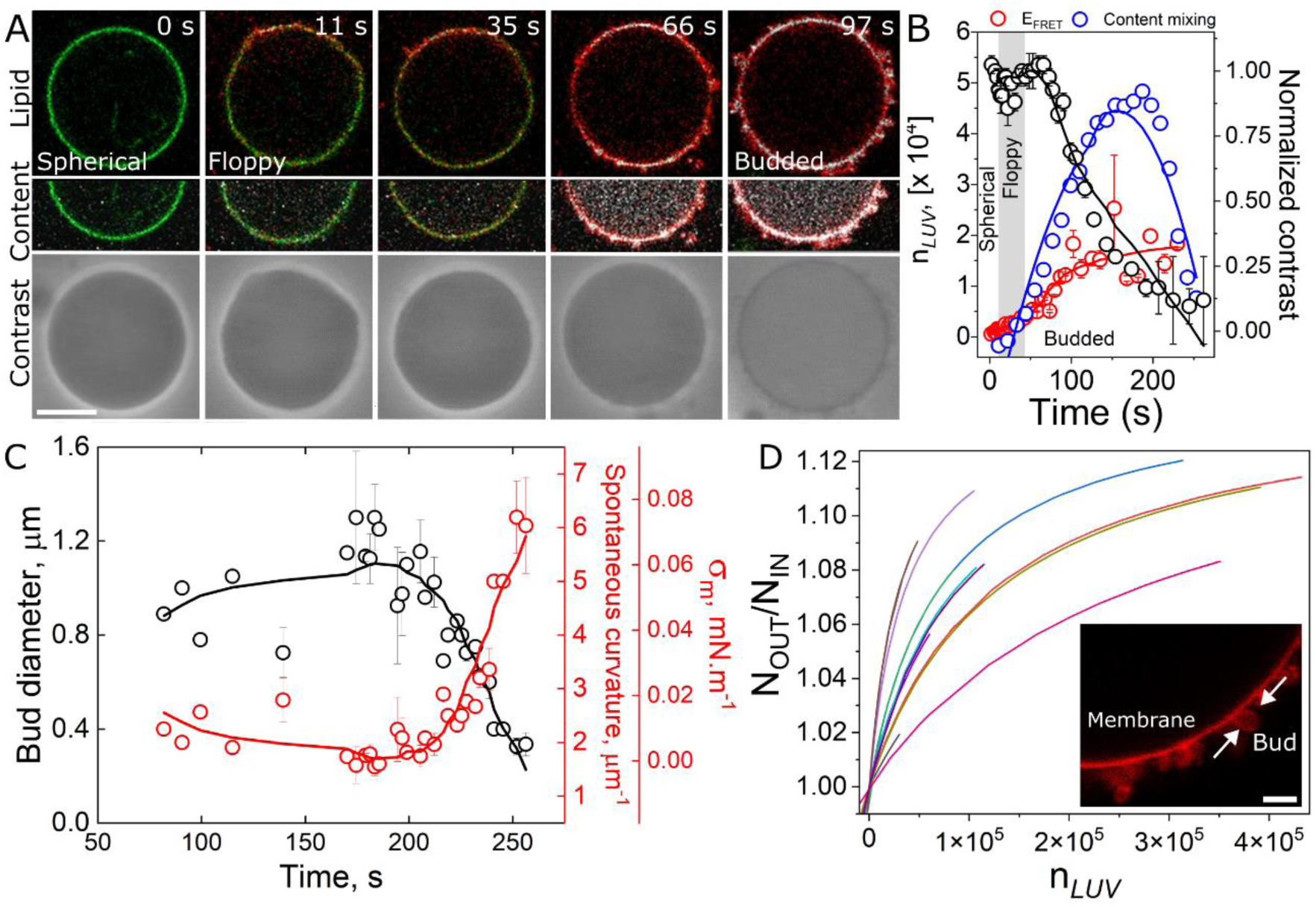
Fusion-driven buildup of membrane spontaneous curvature and tension. (A) Time-lapse snapshots of a representative GUV during fusion with LUVs. The LUVs are injected from above (i.e. the upper part of the image) using a micropipette (not shown). Top row: FRET-based lipid mixing (donor and acceptor excitation/emission). Middle row: Simultaneous monitoring of lipid (green/red) and content (gray) mixing. Bottom row: Phase contrast microscopy showing vesicle morphology. Time stamps indicate seconds from the onset of imaging. Scale bar: 10 µm. (B) Quantification of the number of fused LUVs (*n*_*LUV*_) based on lipid mixing (red) and content mixing (blue), plotted over time. Normalized phase contrast (black) indicates vesicle contrast changes. Annotations reflect key morphological states of the GUV. (C) Temporal evolution of optically resolved bud diameter (black, left axis), along with the corresponding calculated spontaneous curvature *m* and spontaneous tension *σ*_*m*_ (red, right axis). The orange shaded region marks the diffraction limit below which buds cannot be resolved optically. (D) Calculated leaflet lipid number asymmetry, (*N*_*out*_/*N*_*in*_)_*LUV*_, as a function of *n*_*LUV*_. Inset: Zoomed-in region of a GUV membrane densely populated with buds. Scale bar: 2 µm.

GUVs, due to their large size, normally contain nearly equal numbers of lipids in both leaflets. However, small vesicles like LUVs do not exhibit this symmetry. Molecular dynamics simulations have shown that tensionless, highly curved lipid vesicles are intrinsically asymmetric, with more lipids in the outer leaflet(56, 57). This asymmetry alters membrane morphology and tension(58) (also experimentally observed(59) and demonstrated in Figure 4), potentially leading to membrane rupture(57). Since no pores form during early fusion stages, the lipid number imbalance cannot relax, and asymmetry builds up (along with increased spontaneous curvature, budding, and tension). For the LUVs used here, the leaflet asymmetry is approximately (*N*_*out*_/*N*_*in*_)_*LUV*_ ≈1.2 (i.e., ∼55% outer leaflet lipids and 45% inner, see Methods for calculation details), while GUV asymmetry (*N*_*out*_/*N*_*in*_)_*GUV*_ increases from <0.02 to ∼0.115 (Figure 4E), sufficient to drive budding and, eventually, even perforating and causing loss of membrane integrity as indeed observed for high LUV concentration. This observation is consistent with simulation studies(57, 60, 61), demonstrating that lipid number density differences between membrane leaflets destabilize the membrane, occasionally leading to transfer of lipids across leaflets, pores and formation of nonbilayer structures.

Fusion-induced membrane asymmetry is even more pronounced in the case of hemifusion, particularly with neutral POPC membranes, where lipid transfer is limited to the outer leaflet. For example, a 15 μm-radius GUV undergoing hemifusion with 625 LUVs can reach a transbilayer asymmetry of ∼1.17, far exceeding that of fully fused negatively charged GUVs at similar fusion levels. This suggests that at early stages of fusion (low n*LUVs*), hemifusion events produce greater mechanical stress and curvature effects, enhancing the likelihood of membrane destabilization, as indeed observed with POPC GUVs (data not shown).

## DISCUSSION

### Fusion-driven growth and its mechanical consequences

During the cell cycle, membrane area must increase (even duplicate) to support growth and division. One physiological mechanism to achieve this expansion is membrane fusion. However, our study shows that the incorporation of membrane material through fusion of small vesicles does not come without physical consequences. Small vesicles such as LUVs and the intra and extracellular vesicles exhibit an intrinsic lipid number asymmetry, with more lipids residing in the outer than the inner leaflet. When large numbers of such vesicles fuse with a GUV, this asymmetry is transferred to the target membrane, altering its spontaneous curvature and increasing tension. Notably, these are the dimensions of several cellular vesicles.

As a result, the GUV undergoes marked morphological changes, including budding, tension buildup, and even poration and rupture (Figures 2-4). These phenomena have remained underappreciated because they only arise after extensive fusion and such high efficiency is typically not achieved in reconstituted systems. By combining real-time imaging, micromanipulation, and quantitative analysis of fusion intermediates at single-vesicle resolution, we were able to observe these transformations in unprecedented detail. Contrary to other reconstituted systems, the fusion kinetics and efficiencies achieved in our system closely resemble those seen in professional secretory cells, providing a physiologically relevant context for studying membrane remodeling.

### Stages of GUV remodeling during fusion

The membrane remodeling of a GUV following vesicle fusion occurs in a series of distinct and sequential stages, as illustrated in Figure 5. The initial GUV membrane is roughly symmetric. As fusion begins, the GUV acquires additional membrane area, resulting in increased thermal fluctuations. At this early stage, leaflet asymmetry remains low. As the number of fused vesicles increases into the tens of thousands, significant transbilayer imbalances develop, and the GUV enters a budding regime where excess membrane area is stored in visible protrusions (Figure 5B). Figure 5C schematically illustrates our simultaneous quantification analysis of lipid and content mixing, as well as membrane structure and permeabilization.

**Figure 5.**
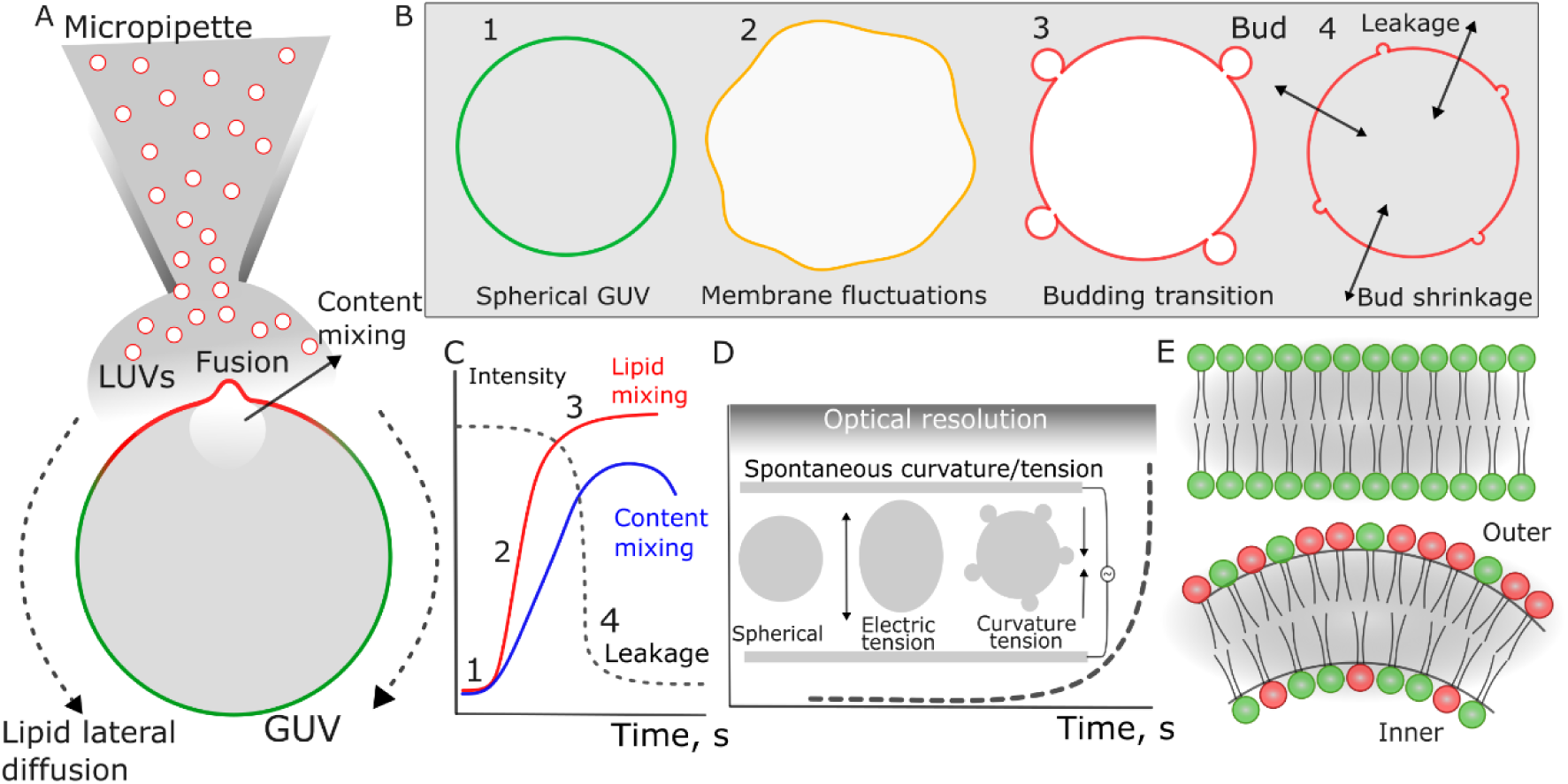
Schematics summary of the observed fusion-induced transformations in GUV morphology and membrane mechanics. (A) Illustration of localized delivery and fusion of LUVs (red), encapsulating a water-soluble content marker (white), to a single GUV (green). Fusion is initiated at the GUV side facing the micropipette, with lipids and content rapidly delivered into the GUV. Lipids incorporated at the fusion site laterally diffuse along the GUV membrane. (B) Sequential GUV morphologies as a function of the total number of fused LUVs, starting from a spherical GUV with a symmetric bilayer (stage 1, see upper sketch in panel E). Progressive fusion leads to membrane area growth and visible fluctuations (stage 2), followed by budding as leaflet imbalances accumulate (stage 3, lower sketch in panel E), and ultimately membrane destabilization perforation when lipid number asymmetry and spontaneous tension becomes critical (stage 4). (C) Readout parameters obtained in real time from the fusion assay, including lipid and content mixing, and membrane permeability, which correspond to the transitions illustrated in B. (D) Diagram showing how fusion-induced changes in spontaneous curvature (dashed line) increase membrane tension, which can eventually exceed the externally applied electric field tension, altering GUV shape during electrodeformation experiments. With increasing spontaneous curvature, bud sizes decrease and can no longer be optically resolves (gray zone). (E) Schematic of lipid number asymmetry resulting from fusion, leading to excess of lipids in the outer leaflet (*N*_*out*_/*N*_*in*_)_*GUV*_ > 1, as a result of lipids transferred from the LUVs (red lipids) to the GUV (green) via fusion.

Upon continued fusion, involving hundreds of thousands of vesicles, the spontaneous curvature further increases (Figure 5D). This leads to the shrinkage of buds beyond optical resolution and drives up membrane tension to the point of leakage and rupture, allowing the contents to escape. At this stage, the asymmetry between the leaflets (*N*_*out*_ /*N*_*in*_)_*GUV*_ can exceed 12%, with a pronounced enrichment of the outer leaflet (Figure 5E).

### Cellular strategies to prevent asymmetry-induced instability

The progression of GUV remodeling events observed here mirrors physical constraints seen in biological systems. For instance, the fusion of around 200 synaptic vesicles per 1 μm² cell membrane at the typical milliseconds rate at the synapse(62), and using 40 nm diameter as the average synaptic vesicle size(63), would lead to a lipid asymmetry ratio (*N*_*out*_/*N*_*in*_)_*GUV*_ ≈1.7 in a few hundreds of milliseconds, which seems an implausible condition. Even more conservative estimates, involving the fusion of 130 vesicles per synaptic button during a period of 10 minutes (64), with the average button area of 1 μm^2^ (65), can already produce noticeable membrane curvature.

Cells have evolved several mechanisms to avoid the pathological consequences of leaflet asymmetry caused by vesicle fusion. First, they regulate the number of fusion events by modulating the abundance of fusion proteins and controlling local membrane properties that influence fusion energetics. Second, they actively remove excess membrane area through fission events, particularly in regions of high curvature. It has recently been shown that area incorporation from as few as 5 single vesicles is sufficient to compress and buckle the acceptor membrane, wherein the excess area is almost instantaneously removed by scission(66).

Another strategy involves the activation of lipid translocation enzymes that redistribute lipids across the bilayer, reducing asymmetry(67). Additionally, cells often maintain membrane reservoirs in the form of nanotubes or invaginations, which serve as mechanical buffers that can absorb excess area without destabilizing the membrane(68). When membrane rupture does occur, dedicated repair machinery is rapidly recruited to reseal the breach and restore homeostasis(69). We have recently shown that fusion of large numbers of LUVs acutely modifies the mechanical properties and structure of cells, such as inducing changes in membrane packing and perforation(36). However, when the vesicles are removed, physiological membrane turnover returns the cells to their steady-state.

The consequences of lipid number asymmetry extend beyond mechanical rupture. Increased spontaneous curvature induces lateral tension, which can activate mechanosensitive ion channels and keep them in the open state under extreme conditions(70), an effect observed across various systems(71). Curved membranes also exhibit altered mechanical properties: they soften, stretch, and become more permeable to ions and small solutes, and they can slow the diffusion of certain membrane proteins(72, 73), a phenomenon attributed to increased membrane tension at high curvatures(74).

To mitigate the consequences of fusion-mediated leaflet asymmetries, cells likely employ several strategies. They may restrict the number of fusing vesicles by regulating the abundance of fusion proteins(75), and locally modulating membrane biophysical properties(34) or the energy landscape of fusion intermediates(76). Second, the incorporation of lipids can reduce tension and cause the acceptor membrane to buckle – particularly in confined regions where asymmetries would accumulate more quickly(77, 78), recruiting curvature-sensing proteins and activating fission to remove excess membrane material(66). Third, transbilayer asymmetries can influence the ability of cell-like vesicles to internalize particles(79), an effect that may have profound consequences in bacterial clearance. Fourth, accumulation of leaflet asymmetries is relaxed by activation of lipid flip-flop machinery(56) or by rapid transbilayer movement of molecules capable of flipping between leaflets(80). Cholesterol, abundant in the plasma membrane, exemplifies such a molecule and has been shown to help balance transbilayer asymmetries(81). Fifth, excess membrane area may be stored as internal membrane invaginations or nanotubes, which can buffer changes in curvature and tension(82) preventing disruption.

### Implications for cell biology, therapeutics, and synthetic systems

It is important to emphasize that the effects we observe, arising from lipid number imbalances across the membrane, are not exclusive to membrane fusion. Large transbilayer asymmetries that increase spontaneous curvature and membrane tension mimic the action of certain antimicrobial peptides(83) and detergents(84, 85), which asymmetrically bind to the outer leaflet and increase membrane tension. Once this tension surpasses a critical threshold, membrane rupture can occur.

Similar physical principles are at play in the endoplasmic reticulum (ER), the site of lipid synthesis in eukaryotic cells. Here, newly synthesized lipids are inserted into the cytosolic leaflet and must be translocated to the luminal leaflet by lipid transport proteins(12). Failure of lipid transport causes leaflet stress, which can lead to budding and eventual loss of organelle function(86). Notably, when membrane area is transferred through hemifusion rather than full fusion, i.e. lipids are added exclusively to one leaflet, the resulting increase in tension is significantly greater. This suggests that hemifusion poses a higher risk of destabilization, and may explain why cells tend to favor docked over hemifused state, explaining why hemifusion is a rather elusive intermediate in cells(87).

In summary, our findings reveal fundamental principles that span cell biology, therapeutics, and synthetic biology. In living cells, the fusion of small, intrinsically asymmetric vesicles is a tightly regulated process, especially in confined regions where asymmetry can rapidly accumulate. In drug delivery, our insights into membrane destabilization mechanisms offer principles for designing fusogenic liposomes that deliver cargo efficiently while minimizing side effects. Finally, in synthetic biology, we highlight the physical constraints imposed by lipid asymmetry during fusion-driven membrane expansion – a key challenge for artificial cell growth and division. By understanding how natural systems manage these constraints, we pave the way for engineering synthetic cells that grow robustly while maintaining membrane integrity.

## CONCLUSIONS

Using a minimal reconstituted system combining live imaging, micromanipulation, and GUVs as synthetic cell compartments, we demonstrated that fusion of small, intrinsically asymmetric vesicles with quasi-symmetric GUVs enables remarkable compartmental growth. At unprecedented rates, reaching several thousand vesicles per second, hundreds of thousands of vesicles can fuse with a single GUV, delivering substantial quantities of both lipid and cargo. However, this dramatic expansion comes with a constraint: the resulting lipid number asymmetries between membrane leaflets limit sustainable growth. The newly incorporated area, initially available for membrane fluctuations and deformation, is rapidly sequestered into highly curved membrane buds as a means of accommodating the imbalance.

In more extreme cases, the spontaneous curvature and mechanical tension induced by asymmetric lipid incorporation compromise membrane integrity, leading to pore formation and leakage of entrapped cargo. The destabilizing effects are exacerbated by smaller vesicles, due to their higher intrinsic asymmetry, and by hemifusion events, in which lipids are transferred into only one leaflet of the bilayer. Notably, these phenomena are not exclusive to membrane fusion but reflect general physical principles likely operating in cellular membranes whenever lipids are asymmetrically incorporated across the membranes.

Our findings have broad implications for cell biophysics, therapeutic delivery, and synthetic biology. Cells appear to use an arsenal of approaches to actively regulate asymmetries, whether they come from fusion or have a different origin. Understanding these regulatory mechanisms can inspire the development of more efficient, less disruptive fusogenic delivery systems. Moreover, insights into how cells manage membrane asymmetry and tension may reveal strategies to overcome key barriers to sustained growth in synthetic cell systems. Harnessing fusion while mitigating its mechanical consequences could pave the way toward building artificial cells capable of continuous expansion and complex functionality.

## MATERIALS AND METHODS

### Materials

The phospholipids 1-palmitoyl-2-oleoyl-sn-glycero-3-phosphocholine (POPC), 1-palmitoyl-2-oleoyl-sn-glycero-3-phospho-(1’-rac-glycerol) (sodium salt) (POPG), 1,2-dioleoyl-sn-glycero-3-phosphoethanolamine (DOPE), and 1,2-dioleoyl-3-trimethylammonium-propane (DOTAP), the fluorescent dye 1,2-dipalmitoyl-sn-glycero-3-phosphoethanolamine-N-(lissamine rhodamine B sulfonyl) (ammonium salt) (DPPE-Rh); and the headgroup-labeled 1,2-dipalmitoyl-sn-glycero-3-phosphoethanolamine-N-(7-nitro-2-1,3-benzoxadiazol-4-yl) (ammonium salt) (DPPE-NBD) were purchased from Avanti Polar Lipids (Alabaster, USA). All lipids were dissolved in chloroform and stored at – 20°C until use. The water-soluble fluorescent dye Atto 647 (Atto) was purchased from ATTO-TEC (Siegen, Germany). Sucrose, glucose and bovine serum albumin (BSA) were purchased from Sigma Aldrich (St. Louis, USA) and used as received.

### Large and giant vesicle preparation

Giant unilamellar vesicles (GUVs) were prepared via electroformation(88) with minor modifications(89). Briefly, ∼ 8 μL of a 3 mM lipid solution containing varying fractions of lipids were spread on conductive ITO-coated coverslips and dried under a stream of N_2_. The glasses were sandwiched using a 1 mm Teflon spacer forming a chamber of ∼ 1.8 mL volume. The chamber was connected to a function generator and an AC field of 1-2 V_pp_ and 10 Hz was applied. The sample was hydrated for 1-2 h at room temperature with a 200 mM sucrose solution in the dark. Although the actual lipid concentration in the GUV preparations are not known, we assume that all lipids used in their preparation resulted in GUV formation, thus yielding a solution of 13.3 μM lipids. GUVs used for the electrodeformation experiments were prepared with a sucrose solution containing 1 mM NaCl. When required, the GUVs were diluted ≈ 5 fold in a slightly hypotonic glucose solution (180-185 mM) to mildly inflate the vesicles and suppress membrane fluctuations. Note that the actual glucose solution in the GUV outside medium is slightly higher as the GUVs were typically diluted ≈ 5 fold.

Fusogenic cationic large unilamellar vesicles (LUVs) were prepared by the extrusion method as previously described(32). In short, a chloroform solution containing DOTAP, DOPE and DPPE-Rh (1:1:0.1 mol ratio) was deposited in a glass vial and the organic solvent was evaporated under a stream of N_2_, forming a film of lipids. Residual traces of solvent were further removed under vacuum for 1-2 h and in the dark. The lipid film was hydrated with a sucrose solution (200 mM) at room temperature at a final lipid concentration of 2.1 mM and vortexed until complete detachment of lipids, forming multilmamellar vesicles (MLVs). In most experiments, the sucrose solution additionally contained 20 μM Atto. The MLVs were extruded with the help of a miniextruder (Avanti Polar Lipid – Alabaster, USA) through 100 nm pore diameter polycarbonate membranes 11 times to form LUVs.

### LUV and GUV incubation

GUVs and LUVs were mixed in different ways: (i) in Eppendorf tubes (Eppendorf AG, Germany) for a specific time and specific LUV concentration, (ii) by injecting LUVs in the vicinity of individual GUVs using a micropipette, or (iii) by hand pipetting LUVs in a chamber with two electrodes in the presence of an AC Field. If not stated otherwise, incubation was performed on slightly hypotonic media to mildly inflate the GUVs and suppress visible membrane fluctuations. In all cases, free Atto present in the medium was not removed but instead it was highly diluted during LUV-GUV incubation (at least 10x depending on LUV concentration). In this case, efficient fusion would result in Atto accumulation in the GUV lumen whose signal is significantly higher than that in the outer medium.

In the first method, increasing concentrations of LUVs (from 0 to 25 μM lipids) were added to a solution containing 30 μL GUVs and 70 μL glucose 180-185 mM. The added GUV volume (and hence concentration) was kept constant so that the only variable is the LUV concentration. The sample was gently and manually stirred and incubated for 10-15 minutes and then placed in a microscope chamber for observation.

In the second method, we prepared micropipettes made from glass capillaries (World Precision Instruments Inc.) that were pulled using a pipette puller (Sutter Instruments, Novato, USA) and tip-forged using a microforge (Narishige, Tokyo, Japan) to obtain tips with an inner diameter around 5 μm. A 420 μM LUV solution (lipid concentration) was loaded in the micropipette and connected to a water reservoir that controls the pressure in the micropipette, and thus the injection flow. LUVs were slowly microinjected a few tens of μm away from individual GUVs and observations were carried out in real time.

In the third method, the GUV solutions (10-50 μL) were diluted in 850 μL glucose 180-185 mM and loaded in an electrofusion chamber (Eppendorf AG, Germany) with parallel cylindrical electrodes (92 μm radius) spaced at 500 μm and connected to a function generator. A weak AC field of 10 V_pp_ nominal voltage and 500 Hz frequency was applied. Afterwards, a small volume (2-10 μL) solution of 0.5-2 mM LUVs was manually injected in the chamber as similar reported for other substances(90). The LUVs reached the GUVs via diffusion.

Prior to all experiments, the chambers were passivated for at least 10 minutes with a 1 wt/volume% BSA and the excess of unbound BSA was removed with excess of glucose. For the microinjection experiments, the micropipette was also passivated with a 1% BSA solution for at least 10 minutes. Very large GUVs (> 60 μm diameter) were not included in the area increase analysis as the fluorescence intensity outside the fusion site did not equilibrate within experimental timescales (∼ 2 minutes).

### Microscopy imaging and analysis

The samples were imaged by fluorescence microscopy using a Leica TCS SP5 or a TCS SP8 confocal microscope (Wetzlar, Germany) using 40X (0.75 NA) air or 63X (1.2 NA) water immersion objectives. DPPE-NBD was excited with an argon laser at 488 nm, and DPPE-Rh and Atto were excited with a diode-pumped solid-state laser at 561 nm and 633 nm respectively, and their emission was detected between 495-555 nm, 565-630 and 640-775 nm, respectively. For continuous real-time recordings, the samples were sequentially imaged with 512×512 pixels, 1 Airy unit, bidirectionally with 3 line averages. The fluorescence of all dyes was detected upon direct excitation and emission. For FRET imaging, DPPE-NBD was excited and emission in the DPPE-Rh channel was detected. These settings were the same for the bulk, micropipette and electrodeformation experiments.

We analyzed the samples by measuring GUV membrane fluorescence in the DPPE-NBD (direct excitation and emission) and FRET channel (direct DPPE-NBD excitation and DPPE-Rh emission). Using LAS X software, a straight line was manually drawn across each GUV and fluorescence intensity along this line was recorded (Figure S1). The maximum fluorescence intensity in each channel was extracted. For the bulk incubation (method i), the fusion signal in the GUV membrane was homogeneous and a line crossing the vesicles on two sides gave rise to two signals, which were averaged and used for the analysis. For methods ii and iii, because the LUVs arrive at the one side of GUVs, the analysis was performed only on the membrane side facing the LUVs as in the example in Figure S3. We used the same software to measure the GUV dimensions, both in the absence and presence of an electric field (which induced GUV deformation). All data were processes and analyzed using OriginPro 16 (OriginLab, Northampton, USA).

## Acknowledgments

The authors acknowledge the MaxSynBio consortium, which is jointly funded by the Federal Ministry of Education and Research of Germany and the Max Planck Society, the E.U. COST action CA22153 ‘European Curvature and Biology Network’ (EuroCurvoBioNet), as well as Fundação de Amparo à Pesquisa do Estado de São Paulo (FAPESP), grant number 2016/13368-4.

Additional information and results are provided in the Supporting information

